# Decision confidence: EEG correlates of confidence in different phases of a decision task

**DOI:** 10.1101/479204

**Authors:** Tanja Krumpe, Peter Gerjets, Wolfgang Rosenstiel, Martin Spüler

## Abstract

Decision making is an essential part of daily life, in which balancing reasons and calculating risks to reach a certain confidence are important to make reasonable choices. To investigate the EEG correlates of confidence during decision making a study involving a forced choice recognition memory task was implemented. Subjects were asked to distinguish old from new pictures and rate their decision with either high or low confidence. Event-related potential (ERP) analysis was performed in four different phases covering all stages of decision making, including the information encoding, retrieval, decision formation, and feedback processing during the recognition task. Additionally, a single trial support-vector machine (SVM) classification was performed on the ERPs of each phase to get a measure of differentiability of the two levels of confidence on a single subject level. It could be shown that the level of decision confidence is significantly reflected in all stages of decision making but most prominently during feedback presentation. The main differences between high and low confidence can be found in the ERPs during feedback presentation after a correct answer, whereas almost no differences can be found in ERPs from feedback to wrong answers. In the feedback phase the two levels of confidence can be separated with a classification accuracy of up to 70 % on average over all subjects, therefore showing potential as a control state in a brain-computer Interface (BCI) application.

## Introduction

Certainty in decision making is an important prerequisite in everyday life, helping to make informed and reasonable decisions in complicated circumstances. Decision confidence plays a role in facilitating adaptive regulation of behavior and supports decision making in complex situations. It affects how subsequent actions are planned or how something can be learned from mistakes that have been made. Decision confidence is also crucial for planning actions in a complex environment especially when subsequent decisions depend on each other or the final outcome of a situation [1,2]. Unfortunately, it is not straightforward to extract decision confidence from behavioral or neurophysiological data as the concept is deeply intertwined with other concepts. One example is evidence or situation evaluation, which is essential to judge a current state correctly. An accumulation of evidence and constant reevaluation of the available facts requires a broad chain of thoughts which interacts with decision confidence [3]. Another example is the strong correlate of reaction time with decision confidence as well as the error rate which also varies greatly with the confidence of the current and previous decisions. It was found that certainty is inversely correlated with reaction time and directly correlated with accuracy and motion strength [4]. Therefore, the level of confidence can easily be confounded with sensory evidence or the planning and execution of motoric actions. In addition, it could be shown that previous choices and the respective feedback influence future decisions [5].

### Perceptual decision making

In the following, we want to steer the focus to perceptual decision making, in which we aim to investigate neural correlates of decision confidence. According to Sternberg and colleagues [6], perceptual decision making can be broken down to three stages: sensory encoding, decision formation, and motor execution. Sensory encoding and decision formation are described in different theoretical frameworks. In a detailed review, Gold and Shadlen made efforts to identify and dissociate the two processes [7]. Two main theoretical groundworks are the basis of this: signal detection theory [8] and sequential sampling framework [9]. Signal detection theory deals with the inability to discriminate between the real sensitivity of subjects and their (potential) response biases caused by conditions of uncertainty. The concept of sensitivity describes the objective difficulty of the task whereas the bias describes the effect of the consequences a decision could have. Missing or detecting a stimulus according to its sensitivity level can be quantified by reaction time and in terms of EEG correlates by P300 latency [10]. The sequential sampling theory, on the other hand, states that the performance of a subject in an experimental task depends on two main factors: the quality of the stimulus information and the quantity of information required before a response is made. More general, Gherman and colleagues [11] state that establishing a certain confidence for a decision relies on the same mechanism as the choice formation itself. Kiani and colleagues [12] found that neurons in the lateral intraparietal cortex (LIP) represent evidence accumulation in monkeys.

### Outcome evaluation

Since it has been established that the outcome of previous choices influences the current ones, we would like to suggest to add a fourth stage to Sternberg and colleagues concept of perceptual decision making, namely outcome and feedback evaluation. As feedback can be very diverse, we want to specify the case we are particularly interested in: Categorical feedback with no further instruction on how to process or use the feedback later in time. Therefore, this goes down to simple performance evaluation and the perception of the latter. General effects concerning the neural correlates following positive or negative feedback that can be found in almost all settings are error-related potentials and feedback-related negativity (FRN). Both belong to the class of event-related potentials (ERPs). Error-related negativity (ERN) for example, was observed in 1990 by Falkenstein et al. [15], time-locked to the presentation of an erroneous event peaking at 80-150 ms. The potential appears strongest at frontal and central electrode sites, has its origin in the anterior cingulate cortex (ACC) [16] and it seems to be linked to error processing [17] and reward prediction [18]. The error-related negativity is often followed by error-related positivity peaking 250-500 ms after stimulus onset which is generated in the posterior cingulate cortex (PCC). This positive component is associated with conscious error perception [19]. Apart from that, there are also event related potentials that are specifically associated with feedback perception. Especially the feedback related negativity (FRN) is a phenomenon often reported, as a negative deflection 145-300 ms after unexpected feedback [20]. It is located frontocentrally and seems to be equal to the N200 component. Interestingly the FN only appears when the feedback is presented immediately after a decision or reaction. The time frame which can still be seen as immediate is at least one second long, according to Weinberg and colleagues [21]. When too much time passes the FN is no longer visible, the P3 component remains unaltered though even if the delay is up to six seconds long. With respect to decision confidence, it was reported that error-related EEG signals vary in a graded way with the level of confidence [13] and adding to that it was found that error positivity (Pe) varies in amplitude with subjective confidence. Both facts show that decision confidence and error detection are closely related processes [14].

### Aim of this study

The aim of this study was to investigate in which stages of perceptual decision making, correlates of decision confidence or certainty can be found in EEG signals. Our interest was to uncover the basic processes of confidence. Therefore, a task design was chosen that allows investigating all four stages of decision making including the process of stimulus encoding followed by decision formation as well as the actual decision making and lastly, the feedback evaluation. The analysis includes classical ERP analysis, as well as machine learning based classification approaches to reveal differences in the EEG correlates between two levels of decision confidence (high and low). In this context, we also evaluated the potential usage of decision confidence as a control state in a Brain-Computer Interface (BCI) application. Having a reliable measure of how confident a subject is during or after a made decision can be a useful information in, for example, educationally oriented applications.

## Materials and methods

A study with two experimental parts was conducted, from here on referred to as part I and part II, to evaluate the impact of decision confidence or other processes that are closely intertwined with the concept of decision confidence. In the following sections the general experimental setup, differences between part I and II as well as the purpose of the differences will be demonstrated. Also, an overview of the used analysis techniques and methods will be given.

### Participants

Part I of the study was conducted on 10 healthy subjects (5 female), with normal or corrected to normal vision. Seven subjects were right-handed and on average the subjects were 22.7 (±3.91) years old. Part II of the study was conducted on 11 healthy subjects (9 female), all right-handed and with normal or corrected to normal vision (age on average 20.45 ± 1.13 years). Due to technical issues, one subject of part I and two subjects from part II were excluded from the analysis, leading to a set of 9 subjects each for both experimental parts. The participation was voluntary and could be ended at any time if required. The subjects received a monetary reward of 8 euro per hour or credits relevant for their study. The study was approved by the local ethics committee of the Eberhardt Karls University of Tübingen and written informed consent was obtained from all participants.

### Apparatus and procedure

#### Technical setup

The subjects were seated in front of a computer screen (19 inches) on which the experiment was presented. The experiment was programmed and presented in Matlab using the cogent graphics extension. A standard keyboard was used for entering the answers by the subject. For the recording of the electroencephalogram (EEG) data, the software BCI2000 [22] was used sampling the data with a frequency of 512 Hz. A Brainproducts Acticap system and two 16 channel g.tec g.USBamp amplifiers were set up for the EEG recording. The integrated high pass filter was set to 0.1 Hz and the integrated low pass filter to 100 Hz. Additionally, a notch filter between 48-52 Hz was applied to eliminate power line noise. 29 electrodes were used for the recording and placed according to the extended 10-20 system (FPz, AFz, F7, F3, FZ, F4, F8, FC3, FCz, FC4, T7, C3, Cz, C4, T8, CP3, CPz, CP4, P7, P3, PZ, P4, P8, O1, Oz, O2, PO7, POz, PO8) and three additional electrodes were used for electrooculogram (EOG) recordings at the outer canthi of the eyes and one on the forehead between the eyes. The ground and reference electrodes were placed on the right and left mastoid respectively and impedances were kept below 10 kΩ. To ensure that stimulus timing is accurately saved in the data, we used the parallel port connected to the EEG amplifier.

#### Task and study design

*Part I:* The study at hand is based on a study originally performed by Woodman and Fukuda [23] which we slightly modified. In general, the experiment was divided into a study phase and in a test phase. In the study phase, the subjects were asked to memorize as many pictures as possible from 500 that were presented. In the test phase, the subjects were presented a mixture of old and new pictures and asked to decide for each picture, if it is already familiar or not. A schematic sketch of the course of the experiment can be seen in Fig 1. All pictures were presented in a block design, where one block consisted of 50 pictures after which a break could be made if needed. The continuation of the experiment was controlled manually by button press by the subject. Each picture presentation can be seen as a separate trial. As stimuli, the same picture dataset as in the Fukuda and Woodmans study was used [24]. The dataset contains pictures displaying daily life objects on a white background. In the test phase, a series of 750 pictures were presented to the subject, again in a block design with a break after 50 pictures each. The 750 pictures consisted of the 500 already presented pictures and 250 new ones in a completely randomized order. The subject was asked to decide after each picture presentation if the picture is new or already familiar. To answer the question one out of four options had to be chosen: 100 % new, 75 % new, 100 % familiar, 75 % familiar. The percentage represents the decision confidence of the given answer. To choose one of the four possible answers the keys A, S,Ö or Ä on a standard German keyboard were used. Both hands were positioned on the keyboard, to handle the key on the right and left side equally fast. The different sides on the keyboard represented the categories familiar or new. To avoid confounds due to handedness of the subject, the sides switched after each block. The switch was indicated before the new block starts and within a block, no changes were made to avoid confusion. Which side represents familiar (right or left) was indicated by circles that appeared next to the picture as soon as an answer was required (1000 ms after stimulus presentation). The circles were blue and yellow, representing the categories familiar and new respectively. A represented high confidence (100 %) and S low confidence (75 %) on the left side, Ö (75%) and Ä (100%) the same on the right side. After the choice was made by the subject, feedback was presented for 1 s indicating if the choice, independent of the certainty, was right or wrong. After feedback presentation, the next picture was presented, and the subject had to decide again about the familiarity of the picture.

**Fig 1.**
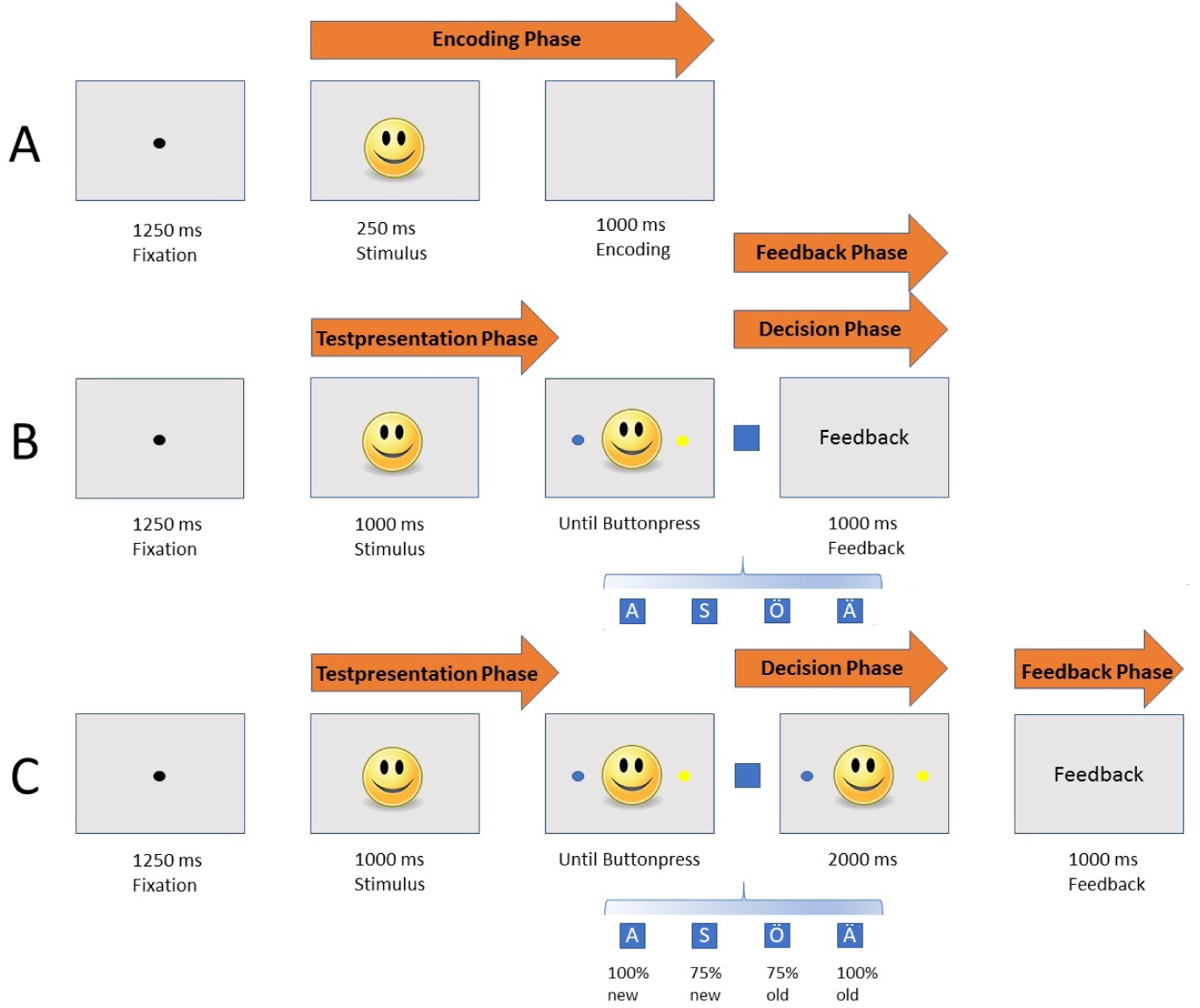
Experimental design. A: Sequence and timing of stimulus presentation during the study phase B: Sequence and timing of stimulus presentation during the recognition test in part I. All phases - depicted by the arrows - are the time frames under investigation. Each time frame accounted for 1250 ms from stimulus onset, except or the feedback phase which was only 1000 ms long. The blue rectangle represents decision via button press C: Sequence and timing of stimulus presentation during the recognition test in part II.

The recording time accounted for 1.15 h on average, from which about 25 mins were needed in the study phase (2.5 s per trial) and about 50 mins in the testing phase (3.23 s per trial plus individual reaction time). The same set of 750 pictures was chosen for each subject, whereas the order of presentation and group affiliation (new or studied) was randomized.

*Part II:* Part II of the experiment differed only in the test phase. 500 instead of 750 pictures were presented. From those 500 pictures, 250 were new and 250 already familiar, therefore the test set of pictures was balanced. Again, the same set of pictures was chosen for each subject, only the order of presentation and the group affiliation (new or old) was randomized. Another difference compared to part I was the timing of the feedback presentation. After making a decision by a button press, a delay of 2 s was introduced before the feedback was presented to the subject. Within the 2 s delay, the stimulus remained on the screen. Despite the reduced number of presented pictures, the duration of the experiment remained almost equal since the individual trials in the test phase were two seconds longer than in part I. Changes were made, on the one hand, to be able to disentangle the decision-making process from feedback processing. A time locked representation of the decision making can only be realized by using the button press as a reference. Since in part I the button press is immediately followed by feedback presentation any correlates related to the decision making might get lost due to new input processing. On the other hand, we wanted to ensure, that no effects due to unbalanced stimuli were introduced in the data, therefore the number of presented stimuli was adapted in the test phase.

### Preprocessing and ERP analysis of the data

The data preprocessing and analysis was performed in Matlab 2015b [25]. Firstly, a bandpass filter between 1 and 40 Hz was applied on the recorded EEG signal and the signal was corrected for EOG artifacts using a regression method proposed by Schoegl [26]. The data was baseline corrected (−100 ms to 0 ms prestimulus or relative to the corresponding event) and cut into trials of one or 1.25 s length depending on the respective categories:

- study presentation phase /Encoding (onset train stimulus presentation 0 - 1250 ms)
- test presentation phase (onset test stimulus presentation 0 - 1250 ms)
- decision phase (onset button press for decision −250 - 1000 ms)
- feedback phase (onset feedback presentation 0 - 1000 ms)

The categorization was chosen to cover all relevant time slots related to the different stages of decision making including the evaluation of the made decision. For each of the categories a further division was made into correct (100 % sure, 75 % sure) and wrong answers (100 % sure and 75 % sure). The four chosen phases will be investigated in terms of behavioral data and ERPs. For the ERP analysis, an additional filtering was performed to exclude trials exceeding 80 or −80 *µ*V from the analysis. The remaining trials of all subjects were averaged for each category individually to reveal differences in the time domain, for high and low confidence answers. After choosing Cz and Pz as representative channels for the evaluation, the statistical significance of the differences in ERPs was established by using a Wilcoxon Ranksum Test [27] on the accumulated signal of all subjects of the respective categories. The resulting p-values were Bonferroni-Holm corrected according to the number of used observations [28]. The significance level was determined to be at p < 0.05. The main interest of the analysis was to extract differences between two levels of confidence (100 % and 75 %) in all four categories. To test for statistically significant differences between the RTs a two-sample t-test was performed.

### Classification

As a second step of EEG analysis, besides the conventional group-based statistics, we used a machine learning based classification approach. This approach is single subject and single trial based, which stands in contrast to grand averages that are calculated over all available trials and subjects. On the one hand, it complements standard analysis, since the single subject information might reveal properties of the data, that the group average can possibly not provide. On the other hand, it is a standard approach that is commonly used in Brain-Computer Interface (BCI) research. In this approach, the data of each subject individually is used to evaluate if the EEG data of the two conditions differs significantly. The machine learning algorithm tries to learn the properties in the data that makes the two experimental conditions distinguishable, which can then be used to decide for each trial to which experimental condition it belongs. This is done by taking all electrodes and therefore, the full spatial pattern of the signal into account. In our approach, a support vector machine (SVM) with a linear kernel (C = 1) was used, as the ML algorithm of choice. The LibSVM implementation [29] for Matlab was utilized in our analysis. The data of 21 channels was used (*3, *z, *4 positions) and the ERP of the phase of interest (1 or 1.25 s time frame) was considered. To prevent over-fitting, a 10-fold cross-validation was performed for each subject and classification. In the cross-validation, the data was divided into ten parts of equal size. In each step of the cross-validation, 90 % of the data are used for training and the remaining 10% are used for testing and evaluating the accuracy of the SVM. In total 10 repetitions are performed in a way that all parts of the data have been used for testing once. The average of all 10 runs is reported as the accuracy for the subject. In all cases, the classes were balanced in size, for training and testing the classifier, to avoid artificial biasing. Additionally, canonical correlation analysis (CCA) was used to generate spatial filters which improve the signal-to-noise ratio of the EEG signal [30]. The filter is calculated on the train data and applied on the test data within each step of the cross-validation. To evaluate the performance of the classification approach, the accuracy was reported, averaged over all subjects. To evaluate the potential influence of the reaction time (RT), the classification was performed on the RT as well. Since a certain level of accuracy can already be reached by chance, depending on the number of classes and used trials per class, the statistical significance of the classification results needs to be established. In order to achieve that we used an approach that estimates the chance level of classification performance by calculating the binomial cumulative distribution [31]. This approach gives rather generalized and conservative bounds, based on sample size and the number of classes. Classification results exceeding the estimated chance level can, therefore, be seen as statistically significant.

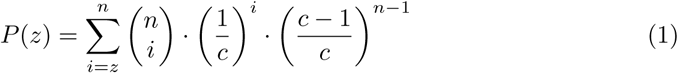

Equation 1 describes the binomial cumulative distribution, in which P(z) represents the probability to predict the correct class at least z times by chance. An appropriate z can be chosen by multiplying the number of samples n with the desired significance level (chosen to be at 0.05). C represents the number of classes and n the number of samples within a class. The approach should only be applied when the classes are balanced. Since they are in our classification approach this is a suitable measure.

## Results

### Behavioral data

#### Accuracy

Table 1 shows the behavioral data that was collected throughout the experiment, representing the number of correct and wrong answers, split according to decision confidence and known and unknown pictures. Presented is the averaged percentage over all subjects for both experimental parts. It can be seen that the proportion of correct and wrong answers is very similar in part I and II. More than 60 % of all pictures have been categorized correctly as new or already familiar and less than 40 % of the trials have been answered wrong. Overall, slightly more known pictures have been identified correctly than new pictures, but there is no major difference between those two categories. In both experimental parts, more wrong answers were given with 75 % confidence than with 100 % across all categories. When comparing the proportions of the correct answers between the levels of confidence it can be seen that there is a major difference between part I and II. In part II the proportion of correct answers is almost equal between the two levels of confidence, whereas in part I more than twice as much correct answers have been given with 100 % confidence compared to 75 %. Despite this seemingly big difference, none of the comparisons showed statistical significance.

**Table 1.** Numbers of correct and wrongly answered trials averaged over all subjects and presented in percent. The results are split according to the familiarity of the pictures (known/unknown) and decision confidence (100/75%).

| # Pictures [%] | All |  |  | Known |  |  | Unknown |  |  |
| --- | --- | --- | --- | --- | --- | --- | --- | --- | --- |
| Part I | All | 100 % | 75 % | All | 100 % | 75 % | All | 100 % | 75 % |
| Correct | 63.6 | 44.27 | 19.33 | 74.40 | 56.40 | 18.00 | 58.00 | 40.00 | 18.00 |
| Wrong | 36.4 | 16.00 | 20.4 | 25.60 | 10.60 | 15.00 | 42.00 | 20.00 | 22.00 |
| Part II |  |  |  |  |  |  |  |  |  |
| Correct | 66.60 | 33.80 | 32.80 | 72.80 | 38.00 | 34.80 | 64.40 | 31.40 | 33.00 |
| Wrong | 34.40 | 15.20 | 19.20 | 27.20 | 12.40 | 14.80 | 35.60 | 14.00 | 21.60 |

#### Reaction time

Table 2 shows the reaction times averaged over all subjects again split according to decision confidence, known or unknown pictures and correctness of the given answers for both parts of the experiment. It can be seen that in part II the subjects were consistently slower for wrong than for correct answers (*p* < 0.01, two-sided t-test and Cohen’s *d* = 0.36), as well as significantly slower for 75 % answers then for answers given with 100 % confidence (*p* < 0.0001, two-sided t-test and Cohen’s *d* = 0.84). In part I this does not seem to be the case, as neither of the two comparisons revealed statistical significance.

**Table 2.** Reaction times of correctly and wrongly answered trials sorted according to the familiarity of the pictures (known and unknown) and decision confidence (100/75%) in ms averaged over all subjects.

| ReactionTime [s] | All |  |  | Known |  |  | Unknown |  |  |
| --- | --- | --- | --- | --- | --- | --- | --- | --- | --- |
| Part I | All | 100 % | 75 % | All | 100 % | 75 % | All | 100 % | 75 % |
| Correct | 1.142 | 1.161 | 1.107 | 1.213 | 1.226 | 1.182 | 0.968 | 0.938 | 0.998 |
| Wrong | 1.132 | 1.123 | 1.137 | 1.119 | 1.051 | 1.158 | 1.150 | 1.203 | 1.103 |
| Part II |  |  |  |  |  |  |  |  |  |
| Correct | 0.934 | 0.840 | 1.126 | 0.953 | 0.822 | 1.166 | 0.959 | 0.860 | 1.091 |
| Wrong | 1.151 | 1.006 | 1.264 | 1.148 | 0.985 | 1.261 | 1.185 | 1.032 | 1.267 |

### Neurophysiological data

#### Encoding phase

In the encoding phase, the first encounter with the stimuli that need to be memorized takes place. In Fig 2 and 3 the ERPs of the channels Cz and Pz respectively, are displayed for all phases in chronological order of the experiment. It can be seen that the encoding phase looks very similar between the two experimental parts at position Cz, with the exception that in part II some points in time differ significantly between the ERPs of the two levels of confidence, whereas they do not in part I. In general, the ERPs are characterized by a positive peak at 200 ms and another large positive component at around 600 ms. In part I there are two small negative components directly preceding the positive components, which are less distinct but still visible in part II. In both parts, statistically significant differences can be found in the ERPs with respect to the two levels of confidence.

**Fig 2.**
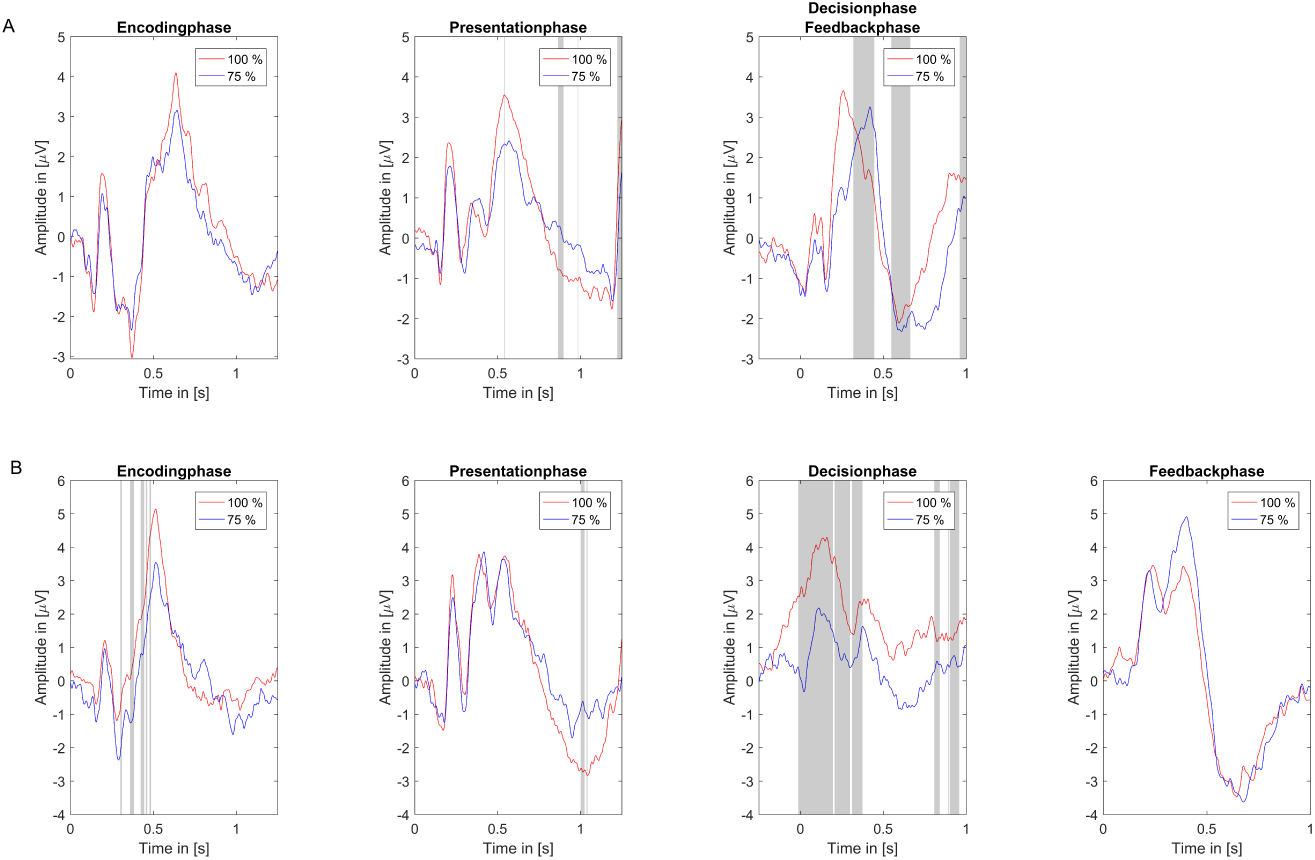
ERPs of confidence levels at Cz. Levels of confidence (100 % red and 75 % blue) for all four phases at electrode position Cz. Significant differences in signal (p < 0.05, Bonferroni-Holm corrected) are indicated by the shaded gray areas. A: Part I (top row), B: Part II (bottom row)

**Fig 3.**
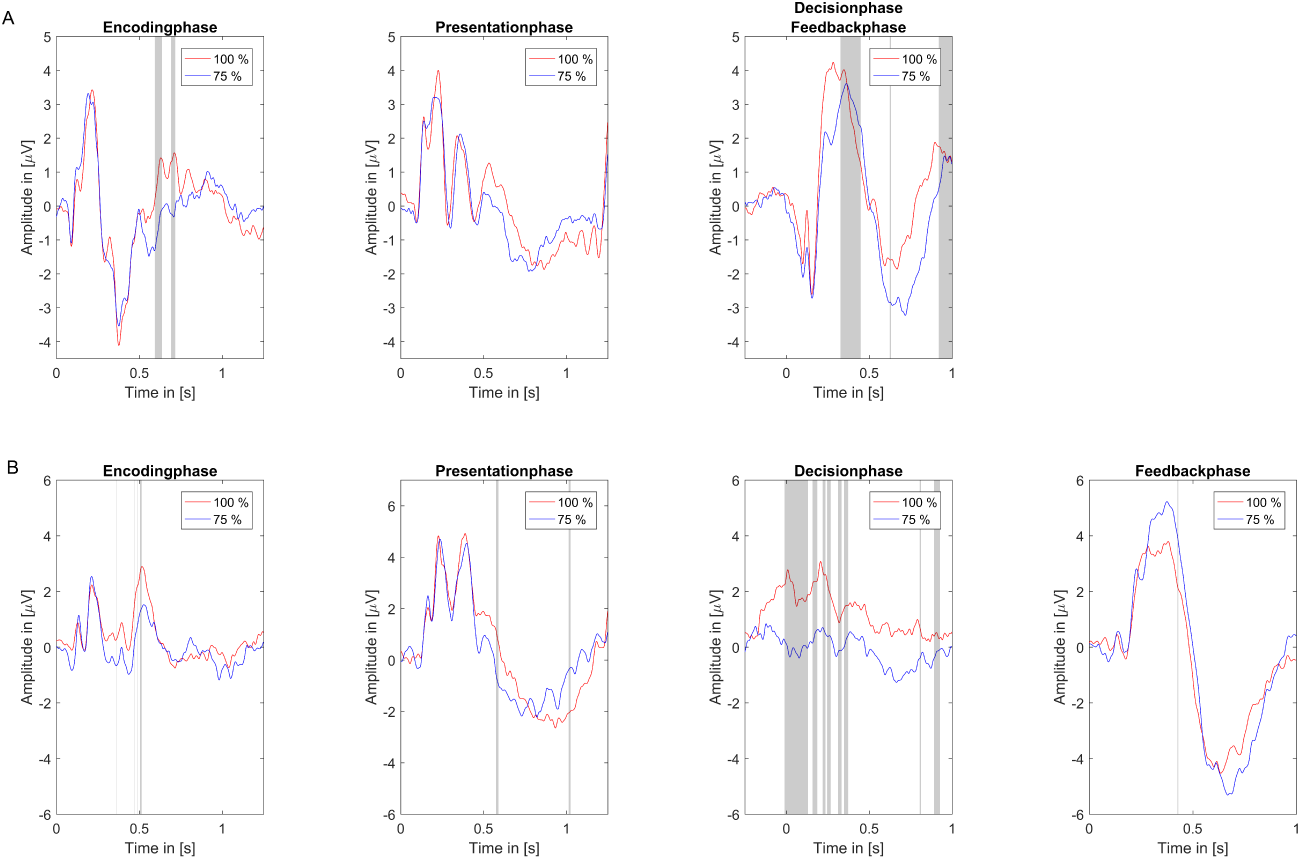
ERPs of confidence levels at Pz. Levels of confidence (100 % red and 75 % blue) for all four phases at electrode position PZ. Significant differences in signal (p < 0.05, Bonferroni-Holm corrected) are indicated by the shaded gray areas. A: Part I (top row), B: Part II (bottom row)

#### Test-presentation phase

The test presentation phase is investigated as a second time frame of interest with respect to the EEG correlates related to decision confidence. Again, the ERPs at positions Cz and Pz can be seen in Fig 2 and 3. At Cz, the ERPs are characterized by two negative components at 150 ms and 300 ms, and three positive components at 200, 400 and 500 ms after stimulus onset. It can be seen that the P400 is much smaller in part I than in part II, which is statistically significant (see Fig 4). The overall differences between 100 and 75 % are rather small and only significant for very few points in time in both parts of the experiment and for both electrode positions.

**Fig 4.**
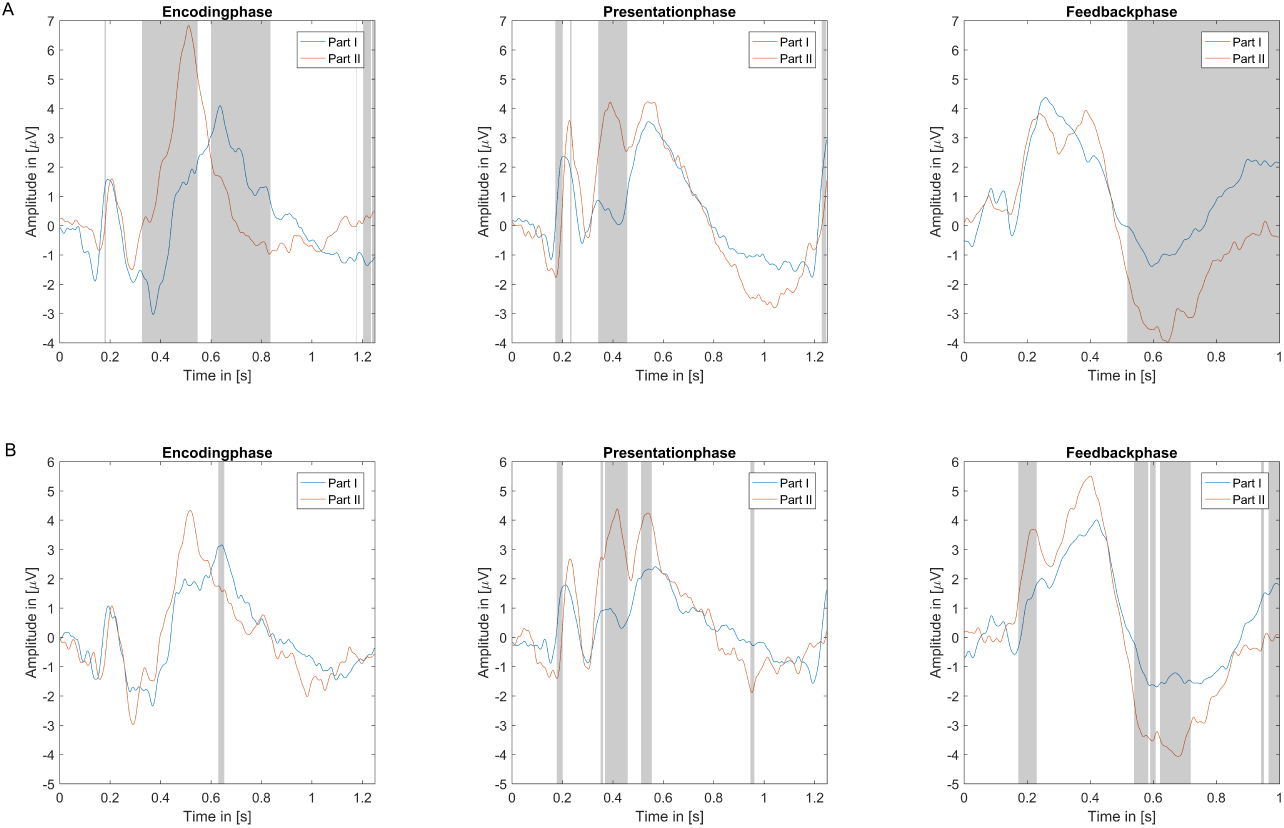
Differences in ERPs between part I and II. Differences between Part I (blue) and Part II (red) for the three comparable phases at electrode position CZ. The shaded gray areas indicate significance (p < 0.05, Bonferroni-Holm corrected) A: 100 %, B: 75 %

#### Decision phase

As a third time frame of interest, the decision phase is investigated. Since this time frame is locked to the button press, which represents the subject’s decision, this phase is intertwined with the feedback phase in part I. In part II a delay was introduced between the button press and feedback presentation to be able to disentangle the two phases. Therefore, the results of the decision phase for part I will be shown in the feedback phase subsection and only the results of part II will be presented here. In the decision phase of part II major differences can be seen between trials answered with 100 % and 75 % confidence at position Cz. They are most distinguishable (statistically significant difference) shortly before and after the button press. The amplitude of trials answered with 100 % confidence is clearly higher during the decision phase as compared to trials answered with 75 % confidence. The ERPs are characterized by two positive peaks at around 150 and 400 ms of which the first peak is much higher than the second one (see Fig 2).

#### Feedback phase

As a last time frame of interest, the feedback phase has been investigated, in which a categorical feedback is presented stating ‘correct’ or ‘wrong’ as a response to the decision made by the subject. In this phase, the reaction to and the evaluation of the feedback can be found. Differences between the levels of confidence can be found in both parts, but also differences between the two experimental parts can be found during the feedback phase. Especially visible and well distinguishable is the level of decision confidence in the feedback phase, defined by the reaction to feedback indicating the given answer was correct. This holds true for both experimental parts. For wrong answers, the difference between 100 % and 75 % is almost nonexistent. In general, the level of decision confidence is well reflected in the feedback phase of part I, by a shift of latency (about 100 ms) in the ERP visible at Cz and a stronger negativity at Pz around 800 ms for answers with 75 % confidence compared to answers given with 100 %. The stronger negativity at Pz is also visible in part II, whereas at Cz no significant difference can be found in part II. When looking at Fig 5 representing the ERPs of correct and wrong answers individually it can be seen, that the potentials of the different categories differ less, for the wrong answers than they do for the correct answers. Especially distinct seems to be the difference between 100% and 75 % within the correctly answered trials. The figure also shows that the neural response to correct and wrong feedback differs significantly. In part II the only difference that remains significant is the difference between 100 % and 75 % confidence of correctly answered trials around 400 ms. When correct and wrong answers are combined no significant difference between 100 and 75 % can be found (see Fig 2). It catches the eye that compared to part I the N200 is not visible at position CZ. Fig 4 shows that at least for the 75 % answers this difference between the two experimental parts is significant. As stated in the previous section, decision and feedback phase are strongly overlapping in part I which is why they are treated as one united phase. The part which refers to the decision making only is restricted to −250 ms to 0 ms before button press. It can be seen (Fig 2 Decision/Feedback phase) that there are no significant differences between the two levels of confidence around the time of button press as it was the case in the decision phase of part II.

**Fig 5.**
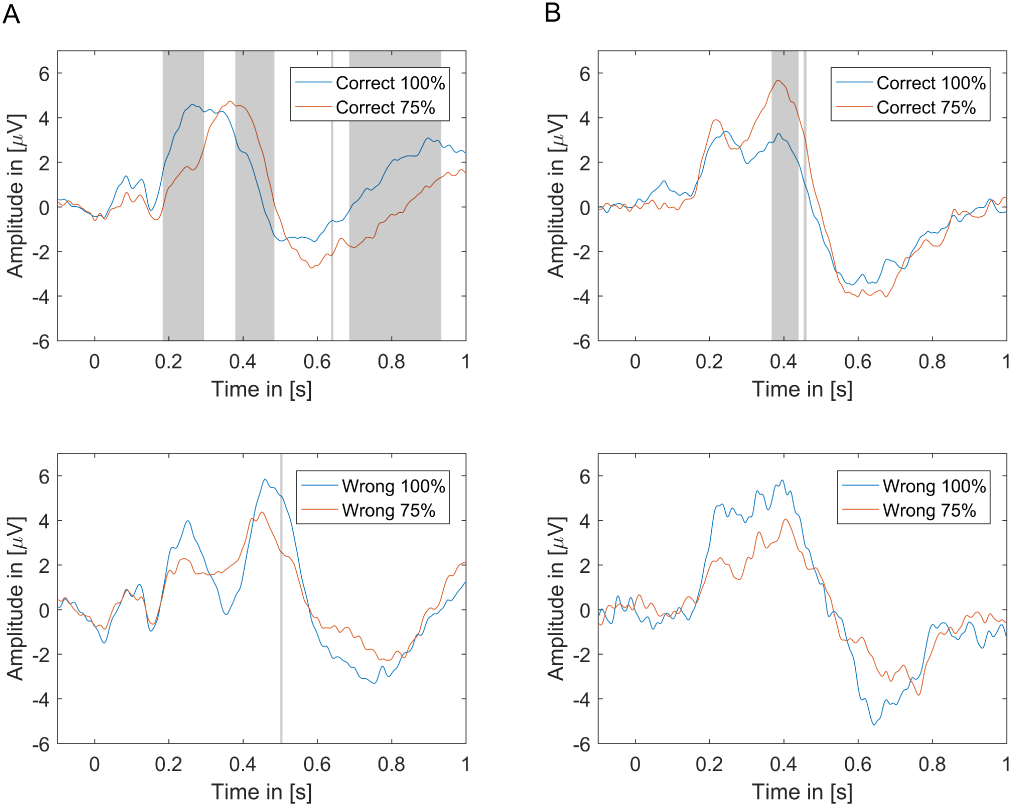
Differences between Part I and II. ERPs at electrode position Cz during the feedback phase. The signal has been split into four categories according to correctness of the answer and the level of confidence. A: Part I (left column) B: Part II (right column). The shaded gray areas indicate statistical significant differences between the categories (p< 0.05, Bonferroni-Holm corrected)

#### Classification

The results of the classification approach, quantifying the success of a single trial separation between the two levels of confidence can be seen in Table 3. It revealed that there are statistically significant differences in all phases for part II and for most phases of part I of the experiment between the ERPs of the two levels of confidences. Fig 6 provides an overview of the accuracy achieved for each subject individually for all phases of the experiment. In addition, it also provides information about the statistical significance as well as the number of available trials in each classification. Despite the statistical significance most of the results do not exceed 60 % accuracy, except for the feedback phase of part I. In addition to using the ERPs of the different phases as features, the reaction time of each trial was also used as a single feature to classify the level of decision confidence. For part II this worked rather well and accuracies above chance level could be reached. In part I the achieved accuracies were not above chance level, indicating that there might be no significant difference. The results for the classification accuracy of wrong trials only are not listed because for many subjects there were not enough trials from this category available to compute the 10-fold cross-validation.

**Table 3.** Classification on ERPs (CCA filtered) of the different phases and the reaction time of the subjects to distinguish 100 % vs 75 % decision confidence. The signal of 21 channels and a 1 s time window were used in a SVM with a linear kernel during a 10-fold cross-validation. Accuracies marked with * are significantly above chance level (0.05) according to binomial cumulative distribution. The results for all answers, as well as for correct answers only have been

|  |  | Encoding | Presentation | Decision | Feedback | Reactiontime |
| --- | --- | --- | --- | --- | --- | --- |
| Part I | All | 51.10 % | 57.97 % * | 69.63 % * |  | 53.16 % |
|  | corr. | 46.08 % | 58.48 % * | 67.64 % * |  | 54.94 % |
| Part II | All | 52.17 % * | 56.24 % * | 59.01 % * | 55.74 % * | 58.38 % * |
|  | corr. | 54.92 % * | 56.16 % * | 58.87 % * | 59.55 % * | 59.52 % * |

**Fig 6.**
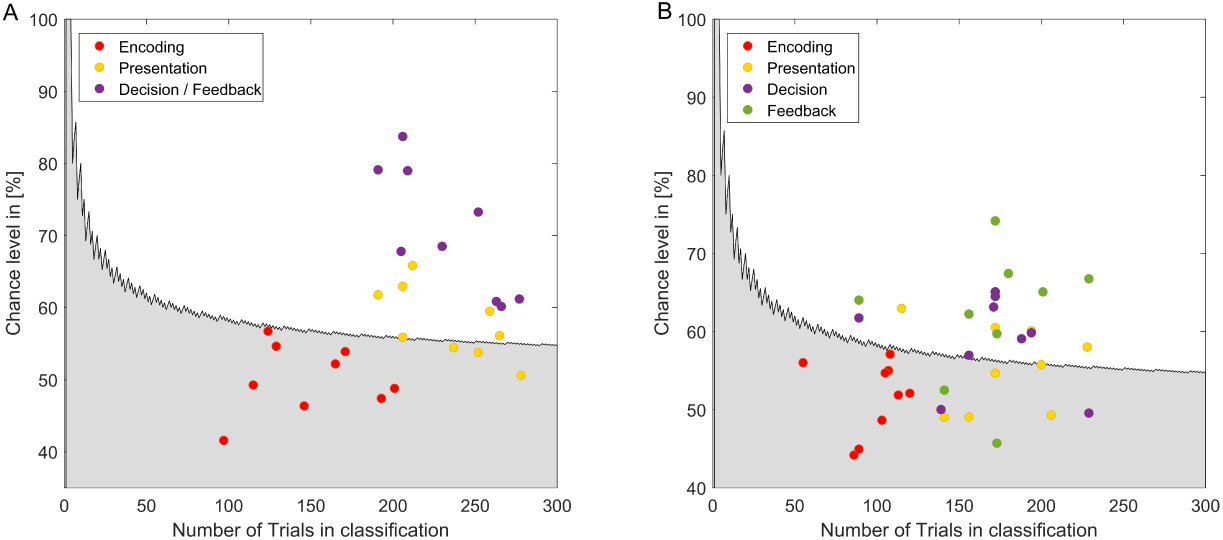
Significance level in relation to the number of trials in all classifications. The shaded area represents all accuracy values that are due to chance in dependency of the available number of trials in classification. As a threshold *p* < 0.05 was chosen. The colored circles show the achieved accuracy values for all subjects 9 for each phase of the experiment. A: Part I B: Part II

## Discussion

In this study, it has been investigated to which degree decision confidence is reflected in ERPs in different phases of a list item recognition task. Although all phases allow a prediction about decision confidence by means of classification accuracy and statistical analysis of the ERPs, the revealed differences might not necessarily represent neural correlates of decision confidence. Nevertheless, we chose decision confidence as an umbrella term, since the labels for all analyzed trials originate from the level of confidence with which the decision was made. To evaluate possible explanations besides decision confidence we will go through all phases step by step. First, we looked at the phase in which the stimulus was encoded in memory. As stimulus encoding is the only process happening at this time point and no decision is involved, there cannot be direct correlates of confidence in this phase. However, attention to the stimulus or the strength of encoding will influence the decision confidence at a later point, and thereby correlates of these processes can be found in the EEG. As the second phase of interest, the stimulus presentation in the test phase was investigated. The test phase started with the instruction to decide about the familiarity of a picture and to state the respective level of confidence. Therefore, in the test presentation phase mainly information retrieval takes place. The decision phase was chosen as an intermediate step to capture the actual process of decision making by choosing a window that starts shortly before the button press (the execution of the decision). Therefore, in this case, it is legit to speak of decision confidence. As a last time window of interest, the feedback phase has been investigated to evaluate if the level of confidence of the decision is also reflected in feedback perception and evaluation. It is possible to speak about affirmation or disappointment which naturally varies with the level of confidence with which the corresponding decision was made. Hence, it can be assumed that at least indirect measures of decision confidence can be measured.

### Behavioral data

In the behavioral data mixed results can be found with respect to decision confidence. The reaction times of part II reflect what can also be found in the literature. The subjects reacted much faster when they were highly confident about their answer, compared to when they were less confident, as well as slower when the answer was wrong than in cases in which the answer was correct [4]. Part I of the experiment does not reflect that. The main reason for that might be the delay of 2 s that was introduced in part II between feedback and decision of the subject. Answering the trials without mandatory breaks, only between blocks, could lead to a loss of focus. Therefore, the subjects needed more time to refocus on a new stimulus and hence also take more time to answer the trial in part I. Another reason could be the shifted proportion of presented stimuli. In part II, the number of known and unknown pictures were equal and in part I it was a ratio of 1 to 3. If the subjects were subconsciously aware of the ratio of known and unknown stimuli remains unclear but it could have an influence on the subjects’ behavior. This fact could also explain the shifted proportions with respect to the level of decision confidence of the given answers. In part I, much more answers have been given with 100 % than with 75 % confidence, whereas in part II the distributions are almost equal between the two levels of confidence. A sort of automatism might have kicked in due to the realization that more known than unknown pictures are presented, resulting in much more high confidence answers.

### Encoding phase

In the encoding phase, some statistically significant differences between trials later categorized with either high or low confidence can be found. Since in the encoding phase no evaluation about the familiarity of the stimulus takes place and therefore no decision needs to be made, the distinguishable levels of confidence do most likely represent other processes. The level of attention paid during the stimulus presentation could be one of them. The more attention has been paid in the encoding phase the easier it is to later categorize the respective stimulus and the more confident will be the answer. Another process that is reflected in the encoding phase could be the actual stimulus encoding. The better a stimulus can be encoded in memory, the easier will be the information retrieval leading to a high confidence answer in the test phase. In the original study of Fukuda and Woodman [23] exactly this process, the quality of memory encoding, was under investigation. The authors showed that the performance of the subjects and the likelihood of high confidence responses were significantly correlated with signal strength (occipital alpha power, frontal positivity) of the encoding period during the individual trials. As Fukuda and Woodman did not evaluate the accuracy of a single-trial prediction, the results presented in this study can be used as a partial estimate. While Fukuda and Woodman grouped their trials into high confidence, low confidence and miss, we only compared high confidence (correct answer with 100 %) and low confidence (correct answer with 75 %) and found that these can be separated with 54.92 % in part II, which is significantly above chance level accuracy. However, for part I classification accuracy was not significantly above chance. Regarding the ERPs in detail, the following can be mentioned. The positive component at 200 ms that is clearly visible in the encoding and test presentation phases of both parts can be associated with stimulus presentation, as the stimulus appeared at that exact point in time and the component appears equally in the encoding as well as in the presentation phase. The encoding phase was perceptually identical in both parts, regarding timing and the presentation of the stimuli. Since the number of known and unknown samples was balanced for the test phase in part II to shorten the experimental time and to avoid effects due to over-representation of one group, fewer trials from the encoding phase can be evaluated because of lacking labels (lacking decision of the subject). In part I all pictures from the encoding phase were represented in the test phase, therefore all were judged with respect to their familiarity and labeled with a level of decision confidence. The pictures not represented in the test phase in part II were not judged, hence they cannot be categorized. This might be a reason for differences in the ERPs of part I and II, since part I includes twice as many samples as part II in the encoding.

### Test Presentation phase

In the test presentation phase the statistically significant differences that can be found are most likely due to information retrieval but also decision formation and therefore, they are at least related to decision confidence. The classification performance has improved in comparison to the encoding phase and remains significantly above chance level, but it is still below 60 %. When comparing the ERPs of both parts, a stronger negativity at around 600 ms at Pz and a little later at Cz for answers given with 75 % confidence compared to answers given with 100 % confidence can be found. Additionally, it can be noticed that the P400 is much higher in part II than it is in part I. This difference could be shown to be statistically significant for both levels of confidence. It could be related to the changed ratio of known and unknown stimuli in the test phase, but another reason could also be the delay of 2 s. Due to using the mandatory break to refocus on the upcoming trials, a higher level of attention and concentration could be present.

### Decision phase

In the decision phase, the process of decision making is captured which is highly influenced by the confidence with which the decision is made. The preparation and the actual motor execution will very likely be reflected in this phase but correlates directly related to the confidence level might be as well. Since decision and feedback processing are in parallel in part I the effects for each process individually cannot be disentangled. To be able to capture ERPs related to the decision making that are not confounded with the processing of new visual input or performance evaluation, a delay of 2 s between the button press and feedback presentation was implemented in part II. To capture correlates that lead to the decision, the phase for the analysis was chosen to start 250 ms before the button press and to end 1 s after it. As the decision phase overlapped with the feedback phase in part I those two effects cannot be separated and therefore, it is difficult to compare those results to part II. Only the time frame −250 ms to 0 ms before the button press are equally present in both parts without the interference of other events. In part I there are no statistically significant differences between the two levels of confidence in this time frame, whereas in part II differences can be found shortly before the button press and also after. Since the effect of feedback presentation starts with button press in part I, there are only 250 ms in which only the decision making is reflected. Visual processing, as well as the evaluation of the presented feedback, might be interfering or interlacing with the neural activity related to decision making, leading to the non-significance in part I. In part II the visual presentation did not change after button press while the subject was in expectancy of feedback. Finding indicators for decision confidence shortly before the actual decision making is in line with the literature [11], which also states that decision confidence and evidence accumulation have the same underlying neural generators in the LIP.

### Feedback phase

During feedback presentation processes like outcome and performance evaluation are taking place. It is easy to imagine that the level of confidence with which a decision was made influences the evaluation to a certain degree. In our experiment, we found the most pronounced differences between the EEG signals of the two levels of decision confidence in the feedback phase of part I. The differences lead to a classification accuracy of up to 70 %. In part II classification accuracy is inferior but still significantly above chance level. When looking at the ERPs it needs to be asked why the difference between the levels of confidence at position Cz is reflected in a shift of latency. The suspicion that the shift could be due to the RT since feedback was always given right after the button press, which in turns is specified by the RT, could be disproven. Classification with the RT as features was shown to be not significantly above chance level. Maybe again the ratio of known and unknown pictures is responsible since the shift in latency could not be observed in part II. Another reason why neither the shift nor other statistically significant differences can be found at Cz during part II of the experiment could be the introduced delay of 2 s. Due to disentanglement, no accumulation of correlates related to decision making and feedback processing can take place. It is possible that especially the accumulation of the two processes in neural correlates leads to the pronounced difference in part I thereby, explaining the absence of this pronounced difference in part II. Another aspect that could explain this phenomenon could be a weakened link between own action and the corresponding feedback, altering the processing and the reaction to the feedback. Interestingly clear indicators for the different levels of confidence can only be found in reactions to correct feedback for both parts of the experiment. This result is in line with literature [32], [33], [34] and therefore, not surprising. Lack of significance between confidence levels in the wrong answers could also be due to the available number of trials, which were a lot less than for correct answers, consistently for all subjects. In general, it remains unclear if the neural responses are that well distinguishable because the subjects were forced to assess their level of confidence with every answer, or if the level of confidence would be reflected as well if there was no need to quantify it after each trial. This fact is hard to revise because the subjective level of confidence needs to be collected somehow to be able to label and categorize the data. Still, since the self-assessment of the current progress in learning is an important marker for deciding when a specific content has been learned sufficiently well, it is an interesting finding.

### Usage for BCI applications

Using machine learning approaches to classify and to distinguish two or more classes of EEG signals is a common approach in brain-computer interface (BCI) research. In BCI research, an accuracy of 70 % is commonly seen as a threshold above which the application of BCI is viable [35]. This value is almost reached for the distinction between levels of decision confidence in the feedback phase in part I. In all other phases the reached accuracy values were statistically significant above chance level but still below 60 %. Having knowledge about the level of decision confidence in a BCI application scenario might be interesting for educational purposes. So far it has been shown that it is possible to assess the amount of load a subject is under and to adapt the difficulty of arithmetic tasks to keep the subject within a comfortable range of load [36], [37]. According to cognitive load theory (CLT) [38], the key to successful learning is to avoid cognitive over- or underload and to keep the learner appropriately challenged. Being able to extract and identify content that is not entirely secured in memory could also be beneficial for the process of learning. This specific content could be recapitulated until the subject reaches a higher confidence during answering the question related to the content. This would be a useful extension to error adaptive learning systems, that would only represent the content that has not been learned at all. Therefore, it can be suggested that using the level of decision confidence during a given feedback might be feasible to use in a BCI based learning application. The fact that accuracy values were lower for part II of the experiment does not interfere with this suggestion, as the main reason for the drop in performance was most likely the introduced delay between given answer and feedback. Since in any kind of application scenario a delayed feedback is usually not desired, because the ability to maintain associations between actions and the resulting rewards is required to measure success or performance, this is not a problem.

## Conclusion

It could successfully be shown that trials labeled according to subjective decision confidence, can be separated with statistical significance in all investigated phases of a simple recognition task. The differentiability of high and low confidence levels could be shown by classical ERP analysis, as well as with a machine learning classification approach. It is possible to distinguish two levels of decision confidence, with up to 70 % classification accuracy, based on the ERPs of the subjects elicited by categorical feedback to the given answer. The main effect resulting in this difference is based on the reaction to positive feedback and not on negative feedback. Since the accuracies are sufficiently high, a usage of this knowledge in BCI applications and research seems feasible. While trying to disentangle feedback processing from decision formation, we found that after introducing a delay of 2 seconds between entering the decision and receiving the corresponding feedback the performance drops immensely. This could either be due to not being able to link the made decision to the corresponding feedback anymore or to the disentanglement of the two phases, revealing that the effect is based on an accumulation of the processes of both phases. Using machine learning as a complementing technique to standard analysis approaches has proven to be helpful to create a profound picture of how different certain mental states are based on their EEG signal.

## Acknowledgments

This experiment was realized using Cogent Graphics developed by John Romaya at the LON at the Wellcome Department of Imaging Neuroscience.” This study was funded by the Leibnitz Science Campus Tübingen ‘Informational Environments’ and further supported by the German Research Council (DFG; SP 1533/2-1) and the Open Access Publishing Fund of the University of Tübingen. Tanja Krumpe is a doctoral student at the LEAD Graduate School & Research Network [GSC1028], funded by the Excellence Initiative of the German federal and state governments. The authors declare no competing financial interests.

